# Morphogenesis of bullet-shaped rabies virus particles requires a functional interplay between the viral matrix protein and ESCRT-I component TSG101

**DOI:** 10.1101/2022.12.16.520694

**Authors:** Yukari Itakura, Koshiro Tabata, Takeshi Saito, Kittiya Intaruck, Nijiho Kawaguchi, Mai Kishimoto, Shiho Torii, Shintaro Kobayashi, Naoto Ito, Michiko Harada, Satoshi Inoue, Ken Maeda, Ayato Takada, William W. Hall, Yasuko Orba, Hirofumi Sawa, Michihito Sasaki

## Abstract

Viral protein assembly and virion budding are tightly regulated to enable the proper formation of progeny virions. At this late stage in the virus life cycle, some enveloped viruses take advantage of the host ESCRT (endosomal sorting complex required for transport) machinery, which contributes to the physiological functions of membrane modulation and abscission. Bullet-shaped viral particles are unique morphological characteristics of rhabdoviruses; however, the involvement of host factors in rhabdovirus infection, and specifically the molecular mechanisms underlying virion formation are not fully understood. In the present study, we used a siRNA screening approach and found that the ESCRT-I component TSG101 contributes to the propagation of rabies virus (RABV). We demonstrated that the matrix protein (M) of RABV interacts with TSG101 via the late-domain containing the PY and YL motifs, which are conserved in various viral proteins. Loss of the YL motif in the RABV M or the downregulation of host TSG101 expression resulted in the intracellular aggregation of viral proteins and abnormal virus particle formation, indicating a defect in the RABV assembly and budding processes. These results indicate that the interaction of the RABV M and TSG101 is pivotal for not only the efficient budding of progeny RABV from infected cells but also for the bullet-shaped virion morphology.

**Importance:** Enveloped-viruses bud from cells with the host lipid bilayer. Generally, the membrane modulation and abscission are mediated by host ESCRT (endosomal sorting complex required for transport) complexes. Some enveloped-viruses utilize their late (L)-domain to interact with ESCRTs, which promotes viral budding. Rhabdoviruses form characteristic bullet-shaped enveloped-virions, but the underlying molecular mechanisms involved remain elusive. Herein, we showed that TSG101, one of ESCRT components, supports rabies virus (RABV) budding and proliferation. TSG101 interacted with RABV matrix protein via L-domain, and the absence of this interaction resulted in intracellular virion accumulation and distortion of the morphology of progeny virions. Our study reveals that virion formation of RABV is highly regulated by TSG101 and the virus matrix protein.

## Introduction

Rabies virus (RABV) is a well-recognized zoonotic virus that causes a fatal neurological disease in mammals. RABV, belonging to the genus *Lyssavirus* of the family *Rhabdoviridae* in the order Mononegavirales, possesses a negative-sense single-stranded RNA genome. This genome encodes five viral proteins: the nucleoprotein (N), phosphoprotein (P), matrix protein (M), glycoprotein (G), and large protein (L)^1^. During the RABV virion assembly stage, the viral RNA genome is surrounded by RABV N together with RABV P and L to form a ribonucleoprotein (RNP) with a firmly organized helical structure. The assembly of the RNP continues with an interaction with the RABV M, which provides a “skeleton structure,” and the RNP is finally enveloped by the cellular lipid bilayer containing the RABV G^2^. The M-condensing RNP structure is essential for the bullet-shaped viral particles that are a characteristic feature of Rhabdoviruses.

In terms of budding and viral particle formation, the importance of the RABV M and G has been highlighted. The RABV G forms membrane microdomains, creates bud sites, and pulls viral particles^3^. RABV M makes a major contribution to the pushing out and pinching off virions as well as promoting efficient virus budding and particle formation, and it binds to RNPs to enable their assembly beneath the cell membrane^2,4–7^. Although G-deficient RABV forms noninfectious but bullet-shaped particles, M-deficient RABV rarely forms infectious virus and produces filamentous particles^2,8^. Therefore, RABV M is an essential component in the formation of the bullet-shaped particle structure of RABV.

Some enveloped viruses hijack cellular factors including the “endosomal sorting complex required for transport” (ESCRT) machinery, to facilitate the budding process. ESCRTs build complexes by sequentially recruiting ESCRT-0, -I, -II, and -III factors during membrane trafficking, e.g., in the multivesicular body sorting pathway^9–11^. The general physiological roles of ESCRTs include the deformation and abscission of lipid membranes, which occur via a process similar to that of virus budding^9,10^. Viral proteins carry consensus amino acid sequences, such as the PY, YL, and PTS/AP motifs, which are called late (L)-domains because they play a role in the recruitment of ESCRT factors at the late stages of the viral life cycle. For example, the human immunodeficiency virus Gag protein and the Marburg virus and Ebola virus VP40 interact with ESCRT-related proteins, such as Alix and TSG101, via their L-domain to promote budding^12–14^. However, little is known about the involvement of ESCRTs in rhabdovirus infection, including RABV infection. Recent proteomic profiling has revealed that ESCRT-related factors, such as chmp4B, HSP40, Alix, TSG101, and chmp2A, are present in purified RABV virions^15^. In addition, RABV M possesses L-domains consisting of the PY and YL motifs, and a previous study revealed that mutations in the PY motif of RABV M led to both the reduced budding efficiency and pathogenicity of RABV in mice^16^. However, the molecular mechanisms underlying the ESCRT-mediated virion assembly and budding in RABV-infected cells has not been clarified.

In the present study, TSG101, a member of the ESCRT-I proteins, was identified as a binding partner of RABV M during the late stage of the virus life cycle. We demonstrate that both the PY and YL motifs of RABV M act as a functional L-domain to enable the interaction with TSG101. RABV propagated in the absence of TSG101 or a recombinant RABV mutant lacking the YL motif failed to bud from the cell membrane and instead formed disrupted virions. Overall, our findings indicate that TSG101 and the L-domain of RABV M are responsible for efficient virus budding and the bullet-shaped morphology of RABV.

## Results

### RNAi screening identifies TSG101 as an ESCRT factor that supports RABV infection

To identify the ESCRT factors involved in RABV growth, ESCRT-knockdown SK-N-SH cells with a custom siRNA library targeting ESCRT factors were infected with luciferase-expressing RABV, and the progeny virus from the cells was quantified using luciferase assays (Fig. 1A, B). Among 25 ESCRTs and their related factors, the knockdown of *TSG101* was most efficient in terms of decreasing the luciferase signal from progeny RABV in the culture supernatants (Fig. 1B). We confirmed that the knockdown of *TSG101* significantly decreased the progeny virus titer compared with that detected following the control-siRNA treatment (Fig. 1C). In addition, the RABV titer was reduced in a concentration-dependent manner by the TSG101 inhibitor ilaprazole^17^ (Fig. 1D). These results reveal for the first time that host TSG101 is a host factor involved in efficient RABV proliferation.

**Figure 1.**
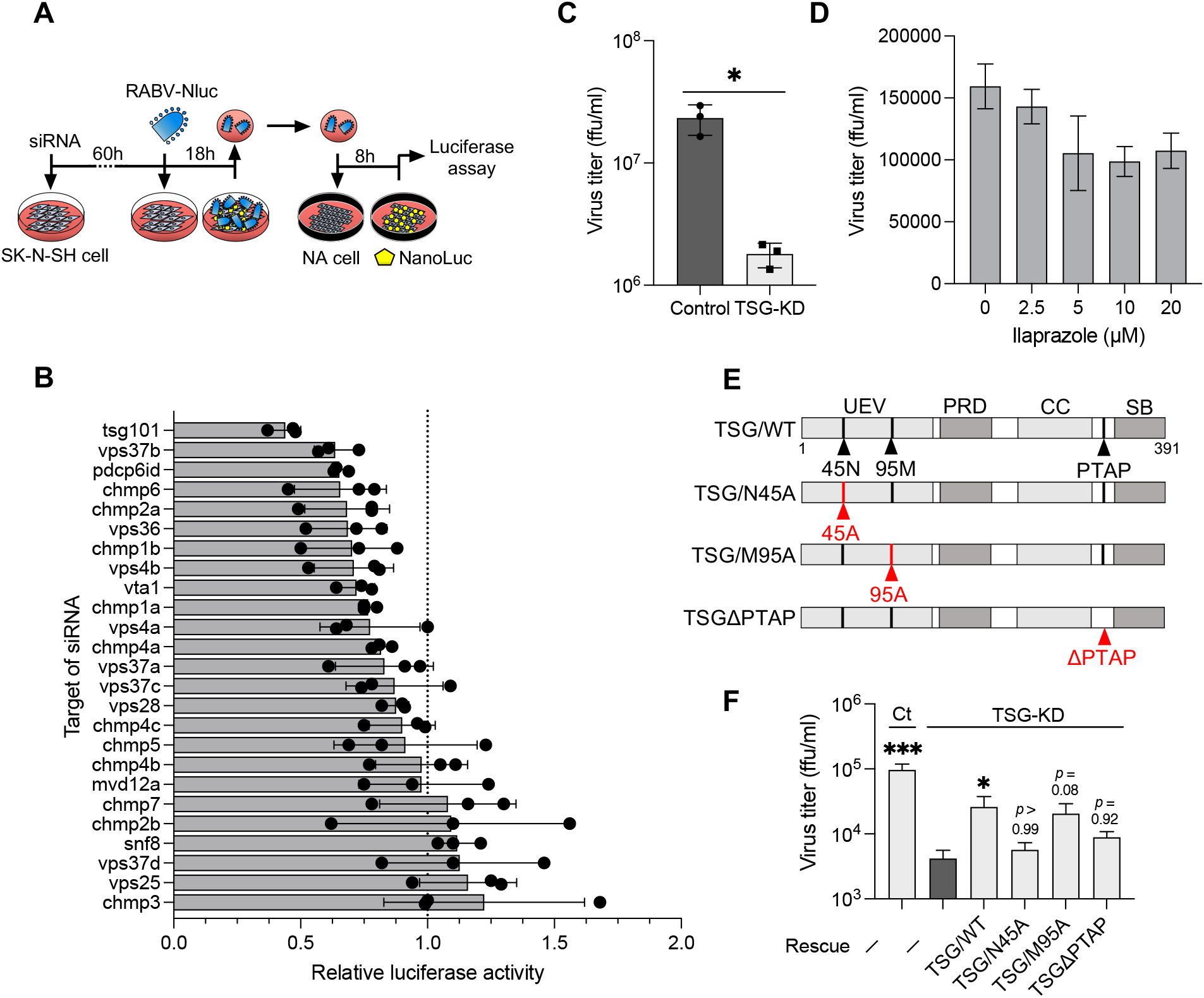
RNAi screening identifies TSG101 as an ESCRT factor supporting RABV infection. (A) Schematic image of the RNAi screening method. siRNA-transfected SK-N-SH cells were infected with RABV-Nluc at a MOI of 10, and culture supernatants collected at 18 hpi were passaged into NA cells. Luciferase activity in the NA cells was measured at 8 hpi. (B) Luciferase activity derived from NanoLuc-encoded reporter RABV in siRNA-treated SK-N-SH cells relative to the luciferase activity in control siRNA-treated cells. Dots indicate the mean of three different siRNAs for each target. Bars indicate the means ± standard deviations of the three siRNAs. (C) Virus titers in the supernatants of TSG-KD SK-N-SH cells at 48 hpi. The titers were measured using a focus forming assay. Bars indicate the means ± standard deviations of three replicates from a representative experiment. (D) Virus titers in the presence of the TSG101 inhibitor ilaprazole. SK-N-SH cells were infected with RABV at a MOI of 1 and cultured with the indicated concentrations of ilaprazole for 24 h. Bars indicate the means ± standard deviations of three replicates from a representative experiment. (E) Schematic images of the TSG101 mutants used in this study. Mutation sites are marked in red. UEV: ubiquitin-conjugating enzyme E2 variant; PRD: proline-rich domain; CC: coiled-coil domain; PTAP: conserved PTAP tetrapeptide motif; SB: steadiness box. (F) Virus titers in TSG-KD and rescue cells. TSG-KD cells were transfected with siRNA-resistant TSG101-encoding plasmids and infected with RABV at a MOI of 1. The virus titers in supernatants at 24 hpi were measured. Bars indicate the means ± standard deviations of three replicates from a representative experiment. Statistical analyses: (C) Welch’s *t*-test: \**P* < 0.05; (F) one-way ANOVA and Dunn’s multiple comparisons tests: \**P* < 0.05, \*\*\**P* < 0.001.

To confirm the TSG101-dependency of RABV infection, we performed a gain-of-function assay by reintroducing TSG101-expressing plasmids into TSG101-knockdown (TSG-KD) cells. In addition to wild-type TSG101, we used three TSG101 mutants: mutants carrying amino acid substitutions at a ubiquitin-binding site (TSG/N45A) or a PT(/S)AP-binding-site (TSG/M95A), and a mutant lacking a PTAP-motif (TSGΔPTAP), which is involved in recognition by the viral L-domain (Fig. 1E). The exogenous expression of wild-type TSG101 significantly rescued the virus titer in TSG-KD cells. Among the three TSG101 mutants, the expression of the TSG/M95A mutant resulted in the highest rescue efficacy, whereas the expression of the TSG/N45A and TSGΔPTAP mutants only exerted partial effects on viral growth in TSG-KD cells (Fig. 1F). These results indicate that the ubiquitin-binding site and PTAP-motif in TSG101 would appear to be required for TSG101-mediated RABV growth.

### Involvement of TSG101 in the assembly and budding processes of RABV virions

Next, we attempted to determine which steps in the RABV life cycle are promoted by TSG101 expression. TSG-KD had no effect on the cellular attachment and entry steps of RABV infection (Fig. 2A, B). A minigenome assay showed that replication and gene expression from a RABV minigenome were not affected by the suppression of TSG101 expression (Fig. 2C). Moreover, intracellular viral RNA levels were similar in control and TSG-KD cells until 24 h postinfection (hpi) (Fig. 2D). Infectious virions were detected from 12 hpi in both cell lines. The virus titer increased markedly at 16 hpi in control cells, whereas a similar increase occurred 4 h later (i.e., 20 hpi) in TSG-KD cells (Fig. 2E). Additionally, the focus size of RABV-infected TSG-KD cells was significantly decreased compared to that of control cells (Fig. 2F), in contrast to the number of RABV-infected foci, which was identical under both conditions (Fig. 2G). The accumulation of RABV M and virions was observed beneath the cell membrane using confocal and electron microscope imaging (Fig. 2H, I). Notably, viral particles released from TSG-KD cells had a rounded form, which is distinct from the typical bullet shape of RABV virions (Fig. 2J). The length of the major axis was 150–200 nm in wild type RABV virions, but the length shifted to 100–150 nm in progeny virions released from TSG-KD cells (Fig. 2K). These results indicate that TSG101 plays a role in the late stage of RABV infection, specifically in virion formation and release as well as subsequent spread to neighboring cells.

**Figure 2.**
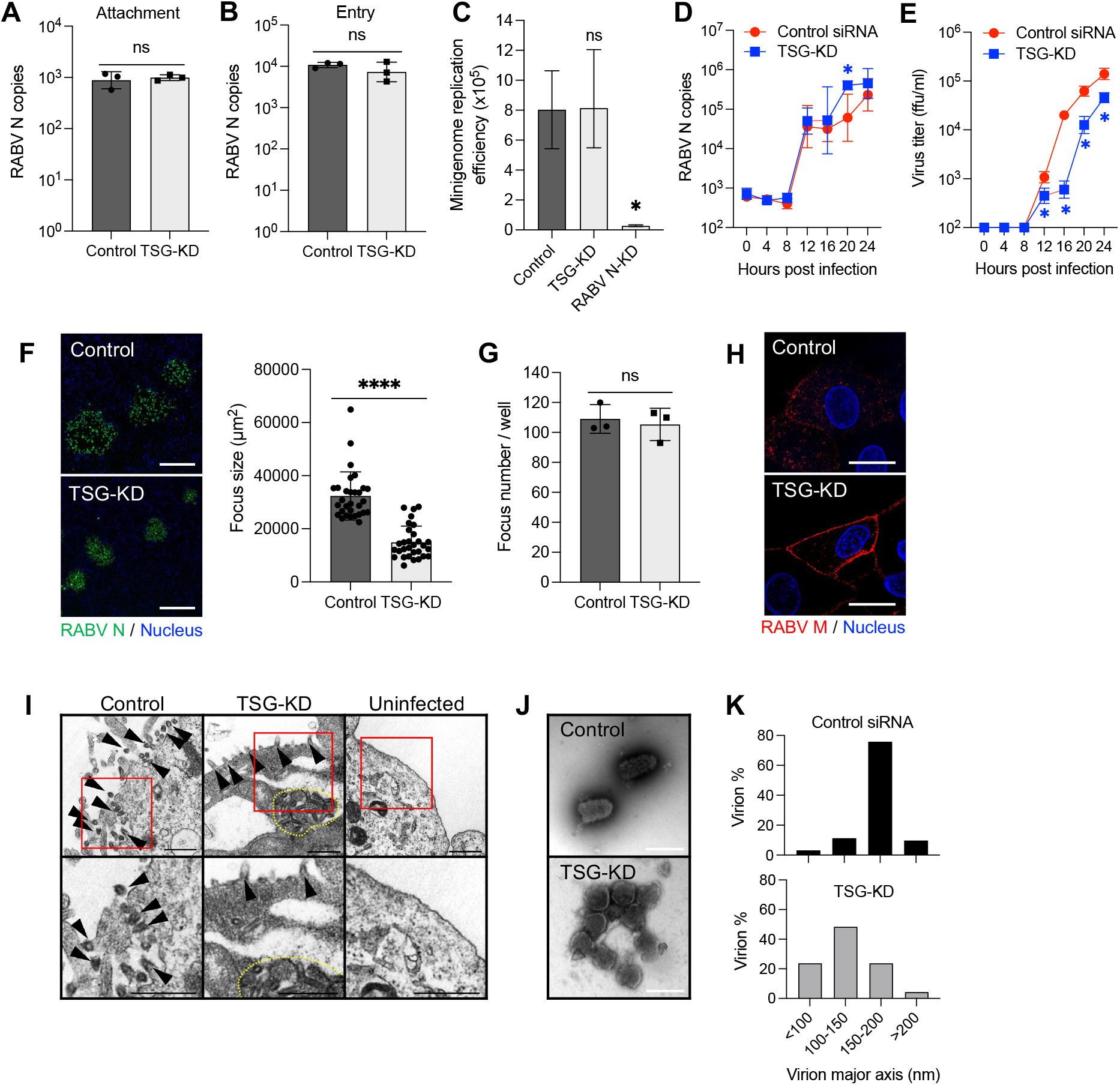
Downregulation of TSG101 expression obstructs the RABV budding process. (A) Virus attachment on the surface of TSG-KD cells. SK-N-SH cells were incubated with RABV at 4L for 1 h. After the cells were washed, RNA was extracted with attached virions and analyzed using qRT-PCR. (B) Viral entry into TSG-KD cells. SK-N-SH cells were infected with RABV at a MOI of 10. After incubation at 37L for 30 min, uninternalized virions were removed via trypsin treatment. Internalized virions were measured using qRT-PCR. (C) RABV minigenome replication in TSG-KD cells. 293T cells exogenically expressing the RABV minigenome were transfected with siRNA against TSG101. Minigenome replication was evaluated by measuring the luminescence signal from NanoLuc. (D) Viral RNA levels at the early stage of virus infection. Viral RNA levels in TSG-KD SK-N-SH cells were measured at the indicated time points using qRT-PCR. (E) Virus titers at the early stage of virus infection. Virus titers in the supernatants from TSG-KD SK-N-SH cells were measured at the indicated time points. (F) Focus size of RABV-infected TSG-KD A549 cells. Foci formed by RABV-infected cells were immunostained with anti-RABV N antibody at 72 hpi. Scale bar: 200 μm. The areas of 30 foci selected randomly were measured using ImageJ. (G) The number of foci in TSG-KD A549 cells. (H) Localization of RABV M in TSG-KD SK-N-SH cells. Cells infected with RABV were immunostained with anti-RABV M antibody at 24 hpi and analyzed using confocal microscopy. Scale bar: 20 μm. (I) Electron microscopic images of RABV-infected TSG-KD SK-N-SH cells at 28 hpi. Arrow heads: virions; dot line: accumulation of virions. Scale bar: 500 nm. (J) Purified RABV virions were negative lystained and analyzed using electron microscopy. Scale bar: 200 nm. (K) Virion diameter and abundance ratio. Purified RABV virions in 50 images captured randomly with an electron microscope were measured using ImageJ. (A–G) Means ± standard deviations of three replicates from a representative experiment. Statistical analyses: (A, B, F, G) Welch’s *t*-test: \**P* < 0.05, \*\*\*\**P* < 0.0001; (C) one-way ANOVA and Dunnett’s multiple comparison test: \**P* < 0.05; (D, E) multiple *t*-tests: \**P* < 0.05.

### TSG101 interacts with the L-domain in RABV M

TSG101 has been reported to interact with the L-domain in the viral proteins of some envelope viruses^12,14,18,19^. RABV possesses two representative L-domains: PY (PPEY) and YL (YVPL) motifs at amino acid positions 35–38 and 38–41 in the RABV M, respectively (Fig. 3A). Therefore, we examined the interaction between RABV M and TSG101. In SK-N-SH cells expressing the TSG101–venus fusion protein, RABV M colocalized with TSG101–venus (Fig. 3B). Additionally, when RABV M and TSG101 were coexpressed in 293T cells, they were also shown to interact in co-immunoprecipitation studies (Fig. 3C). To identify the region responsible for the interaction between RABV M and TSG101, we included RABV M and TSG101 mutants in the co-immunoprecipitation assays. RABV M mutants with a substitution in each L-domain (Fig. 3A) showed lower binding affinity to TSG101 (Fig. 3C). In particular, the lack of the YL motif in RABV M led to a marked decrease in the interaction between RABV M and TSG101 (Fig. 3C). Additionally, TSG/N45A (the ubiquitin-binding deficient mutant) showed a lower interaction with RABV M (Fig. 3C). These results suggest that RABV M interacts with TSG101, and their respective L-domain and ubiquitin binding site are the likely target sites for the interactions.

**Figure 3.**
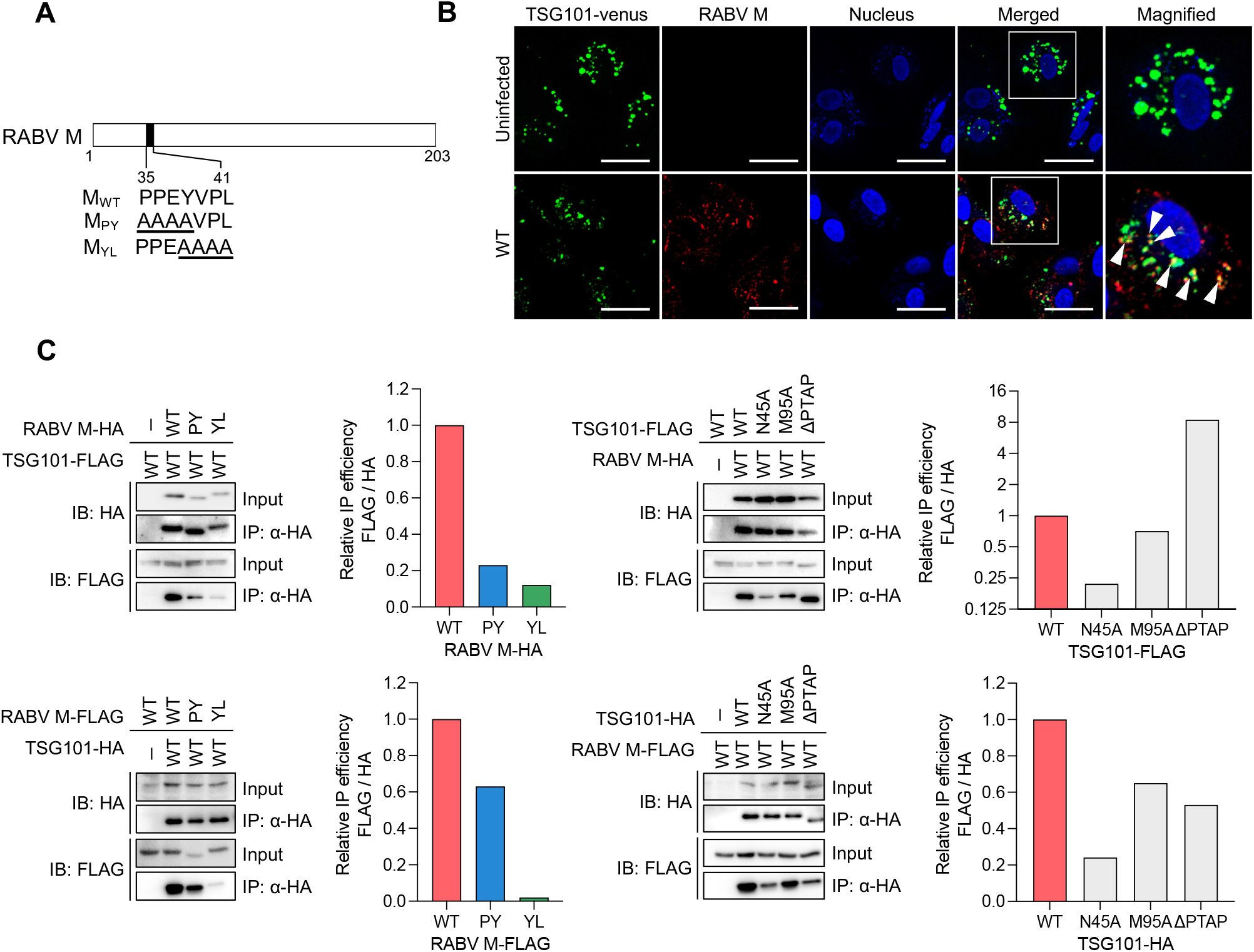
RABV M interacts with TSG101 via the L-domain. (A) Schematic representation of RABV M and the L-domain mutants. (B) Colocalization of RABV M and TSG101. SK-N-SH cells stably expressing TSG101–venus were infected with RABV, immunostained with anti-RABV M at 24 hpi, and analyzed using confocal microscopy. Arrow heads show colocalization. Scale bar: 50 μm. (C) Coimmunoprecipitation of RABV M with TSG101. HA- or FLAG-tagged RABV M and TSG101 were coexpressed in 293T cells and coimmunoprecipitated using anti-HA magnetic beads. Immunoblotting was performed with anti-HA or -FLAG antibody. Bar graphs show the relative precipitation efficacy (IP / Input) of FLAG compared with that of HA from a representative experiment following quantification via ImageJ.

Furthermore, the PY and YL motifs in RABV M are conserved throughout the lineages of RABV (Fig. S1A). As well as the fixed virulent challenge virus standard (CVS) strain, the fixed attenuated high egg passage-Flury (HEP) strain and street Toyohashi strain exhibited significantly decreased virus titers in TSG-KD cells (Fig. S1B), suggesting a general function of RABV M–TSG101 interaction in RABV infection.

### Mutation at the L-domain disrupts the TSG101-dependent infection of RABV

To further determine the importance of the RABV L-domain in the virus life cycle, we generated replication-competent RABV mutant clones, namely RABV/PY and RABV/YL, in which the PY and YL motifs in RABV M were substituted with alanine, respectively (Fig. 4A). We assessed the growth of the mutants in nontreated control cells and TSG-KD cells. In control cells, the virus titer of RABV/YL was significantly lower than that of RABV/WT and RABV/PY; however, there was no difference among the virus titers in TSG-KD cells (Fig. 4B). The accumulation of RABV M in the peripheral cytoplasm, which was observed in TSG-KD cells infected with RABV/WT (Fig. 2H), was reproduced in control cells infected with RABV/PY and RABV/YL (Fig. 4C). In addition, RABV/PY M and RABV/YL M did not colocalize with TSG101–venus in the infected cells (Fig. 4C). These results suggest that both the PY and YL motifs in the RABV L-domain could mediate the RABV M–TSG101 interaction during viral infection, and the YL motif is more important in the TSG101-dependent RABV life cycle.

**Figure 4.**
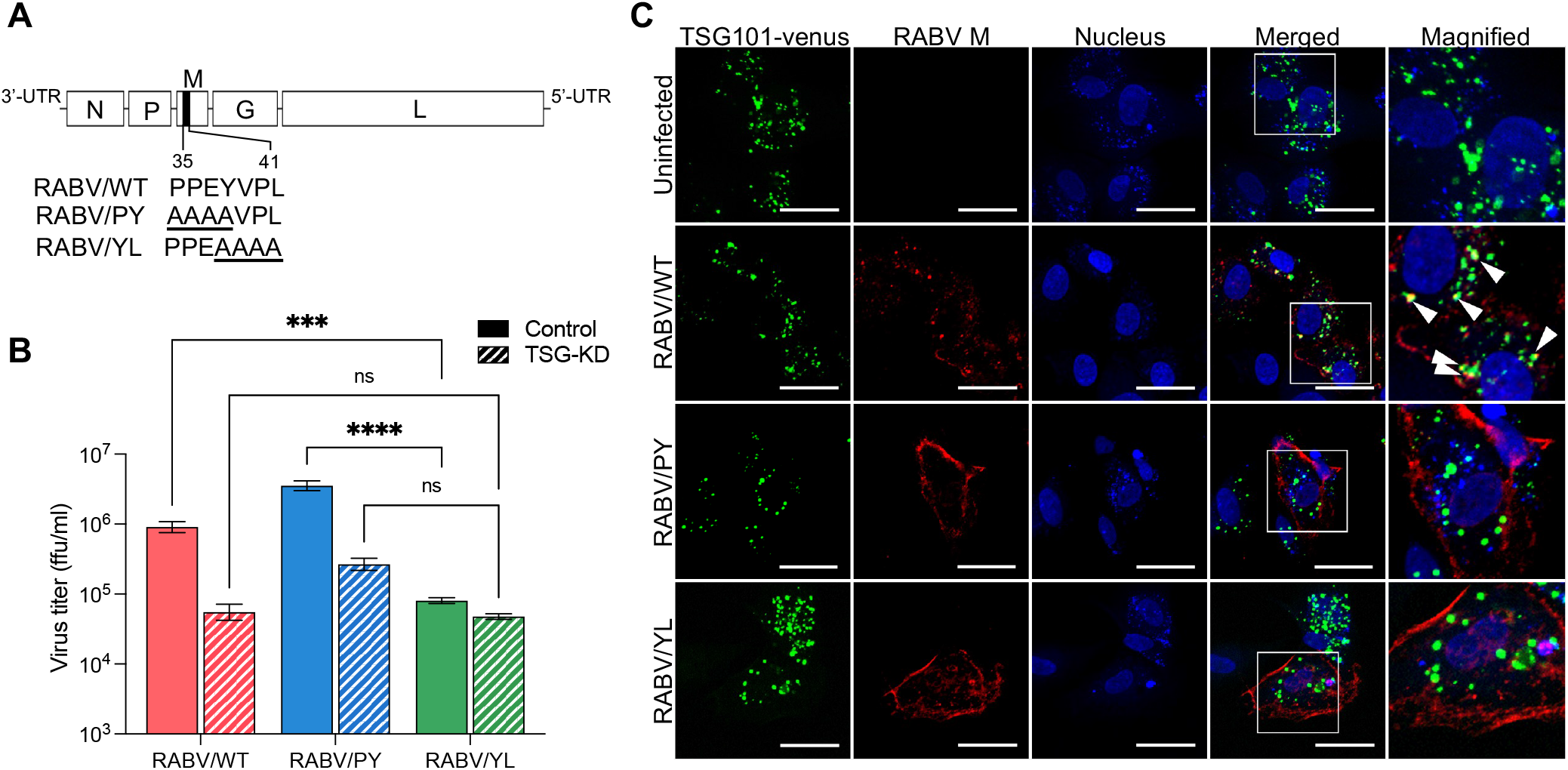
RABV YL motif is essential for TSG101-mediated viral growth. (A) Schematic representation of recombinant RABV with alanine substitutions in the L-domain in RABV M. (B) Virus titers of RABV L-domain mutants. TSG-KD SK-N-SH cells were infected with RABV mutants at a MOI of 1, and virus titers in the supernatants at 48 hpi were measured. Statistical analyses: two-way ANOVA and Sidak’s multiple comparisons tests: \*\*\**P* < 0.001, \*\*\*\**P* < 0.0001. (C) Colocalization of RABV M and TSG101. SK-N-SH cells stably expressing TSG101–venus were infected with RABV, immunostained with anti-RABV M at 24 hpi, and analyzed using confocal microscopy. Arrow heads show colocalization. Scale bar: 50 μm.

### Mutation at the L-domain perturbates RABV budding and spread in vitro

RABV/PY and RABV/YL were further characterized to better understand the role of the L-domain in RABV infection. In SK-N-SH cells, the viral growth of RABV/PY was comparable with that of RABV/WT, whereas the growth efficiency of RABV/YL was lower than that of RABV/PY and RABV/WT (Fig. 5A). RABV/WT and the two mutants showed similar trends in viral RNA replication until 24 hpi (Fig. 5B). Although the infectious progeny viruses of RABV/WT and the mutants were detected in the culture supernatants from 12 hpi, RABV/YL showed a considerably lower virus titer after 16 hpi (Fig. 5C). In terms of focus formation, the focus sizes of RABV/PY and RABV/YL were significantly smaller than that of RABV/WT (Fig. 5D), indicating a low efficiency of virus spread by both mutants. These results indicate that substitutions at the L-domain of RABV M critically decrease viral proliferation. Accompanied by the accumulation of the RABV M along the plasma membrane (Fig. 4C), virus particles remained tethered to the cell surface in RABV/PY and RABV/YL infection, as observed under an electron microscopy (Fig. 5E). Additionally, the accumulation of virions in the cytoplasm of RABV/YL-infected cells was observed. (Fig. 5E). In terms of virion morphology, RABV/PY had slightly longer virions compared with those of RABV/WT, whereas RABV/YL formed rounded particles that clearly differed from the representative bullet-like particles of rhabdoviruses (Fig. 5F, G). Notably, the characteristics of RABV/YL were consistent with those of RABV/WT in TSG-KD cells (Fig. 2D–K). These results suggest that the L-domain, especially the YL motif, in the RABV M plays an essential role in RABV budding and bullet-like virus particle formation.

**Figure 5.**
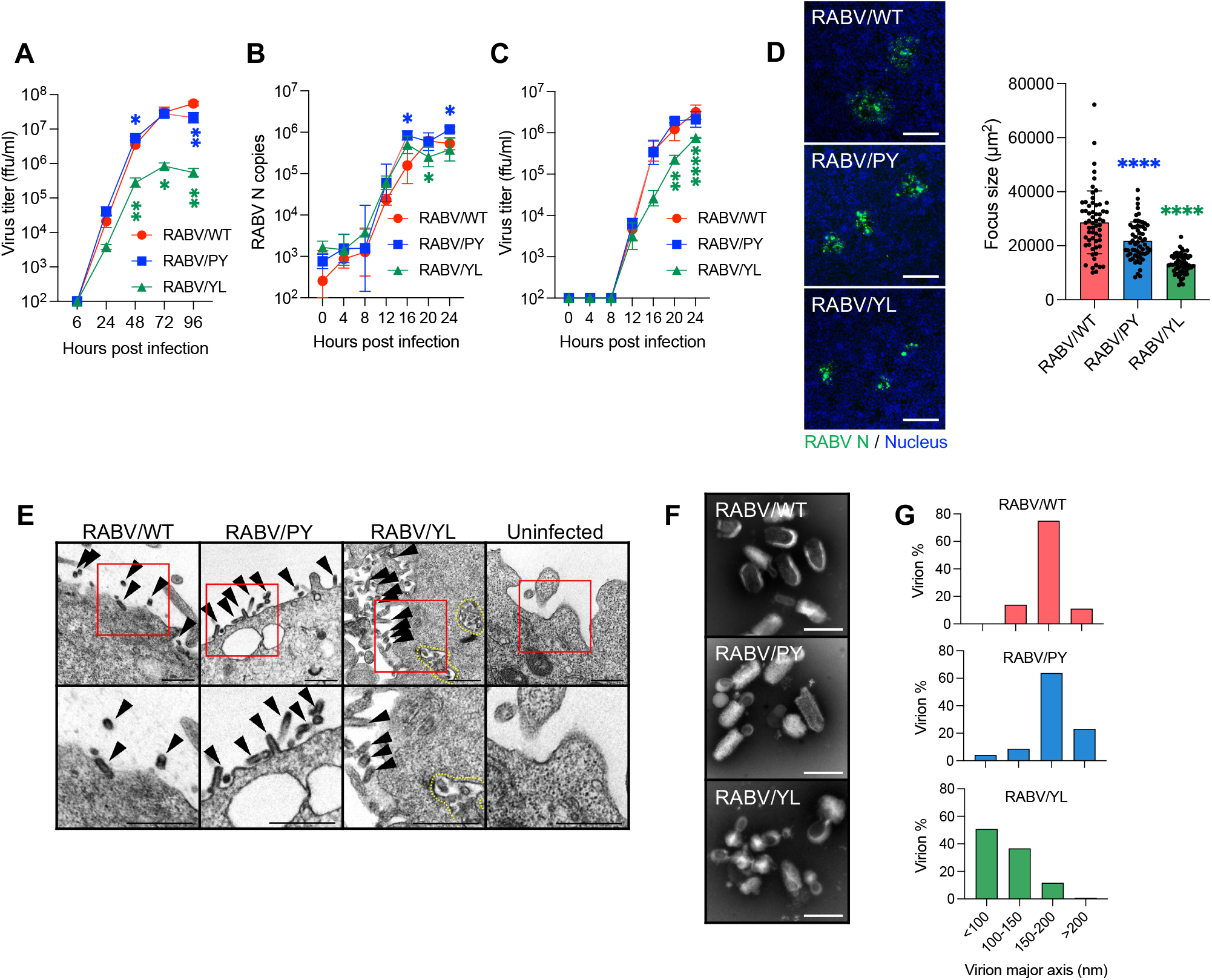
Impaired growth of RABV L-domain mutants. (A) Virus growth curves in SK-N-SH cells during a multicycle infection. Cells were infected with RABV at a MOI of 0.01, and virus titers in the supernatants were measured at the indicated time points. (B) Viral RNA levels at the early stage of virus infection in SK-N-SH cells. Cells were infected with RABV at a MOI of 1, and virus replication in the cells was evaluated at the indicated time points using qRT-PCR. (C) Virus titers at the early stage of virus infection in SK-N-SH cells. Cells were infected with RABV at a MOI of 1, and virus titers in the supernatants were measured at the indicated time points. (D) Focus size of RABV L-domain mutants. The areas of 60 foci selected randomly in NA cells were measured using ImageJ. (E) Electron microscopy images of SK-N-SH cells at 28 hpi of RABV L-domain mutants. Arrow heads: virions; dot line: accumulation of virions. Scale bar: 500 nm. (F) Purified virions of RABV L-domain mutants were negatively stained and analyzed using electron microscopy. (G) Virion diameter and abundance ratio. Purified RABV virions in 50 images captured randomly with an electron microscope were measured using ImageJ. Scale bar: 200 nm. (A–C) Means ± standard deviations of three replicates from a representative experiment. Statistical analyses: (A–C) multiple *t*-tests: \**P* < 0.05, \*\**P* < 0.01, \*\*\*\**P* < 0.0001. (D) One-way ANOVA and Dunnett’s multiple comparison test: \*\*\*\**P* < 0.0001.

### Pathogenicity of RABV L-domain mutants in vivo

To determine the influence of the L-domain on RABV pathogenicity in vivo, mice were intracranially or intramuscularly infected with the RABV L-domain mutants. Intracerebral inoculation showed a 1- and 2-day delay in the survival curve and body weight loss for RABV/PY and RABV/YL mutants, respectively (Fig. 6A, B). The virus titers of both mutants in the brain at 4 days postinfection (dpi) were similar, and they were ∼80-fold lower than that observed with the parental RABV/WT (Fig. 6C). Viral proteins were distributed extensively across the whole brain of RABV/WT-infected mice, whereas infection was limited in the brains of mice infected with RABV/PY and RABV/YL (Fig. 6D, S2). Even on 6 dpi, virus growth and distributions of RABV/PY and RABV/YL were limited compared to those of RABV/WT at 4 dpi, whereas the mice were on the similar disease progressions (Fig. S3, 6C). Following intramuscular inoculation, both mutants had a 2-day delay in the survival curve and body weight change compared with the control (Fig. 6E, F). In terms of virus propagation, the virus titer in the brain was 34.8-fold lower for RABV/PY and 509.6-fold lower for RABV/YL compared with the control virus titer (Fig. 6G). Although the disease progression of mice infected with either mutant was similar, there was a marked difference between the virus titers of the two mutants (Fig. 6G). These results suggest that the substitution of the L-domain results in the attenuation of RABV, indicating the importance of both the PY and YL motifs in RABV replication not only in vitro but also in vivo.

**Figure 6.**
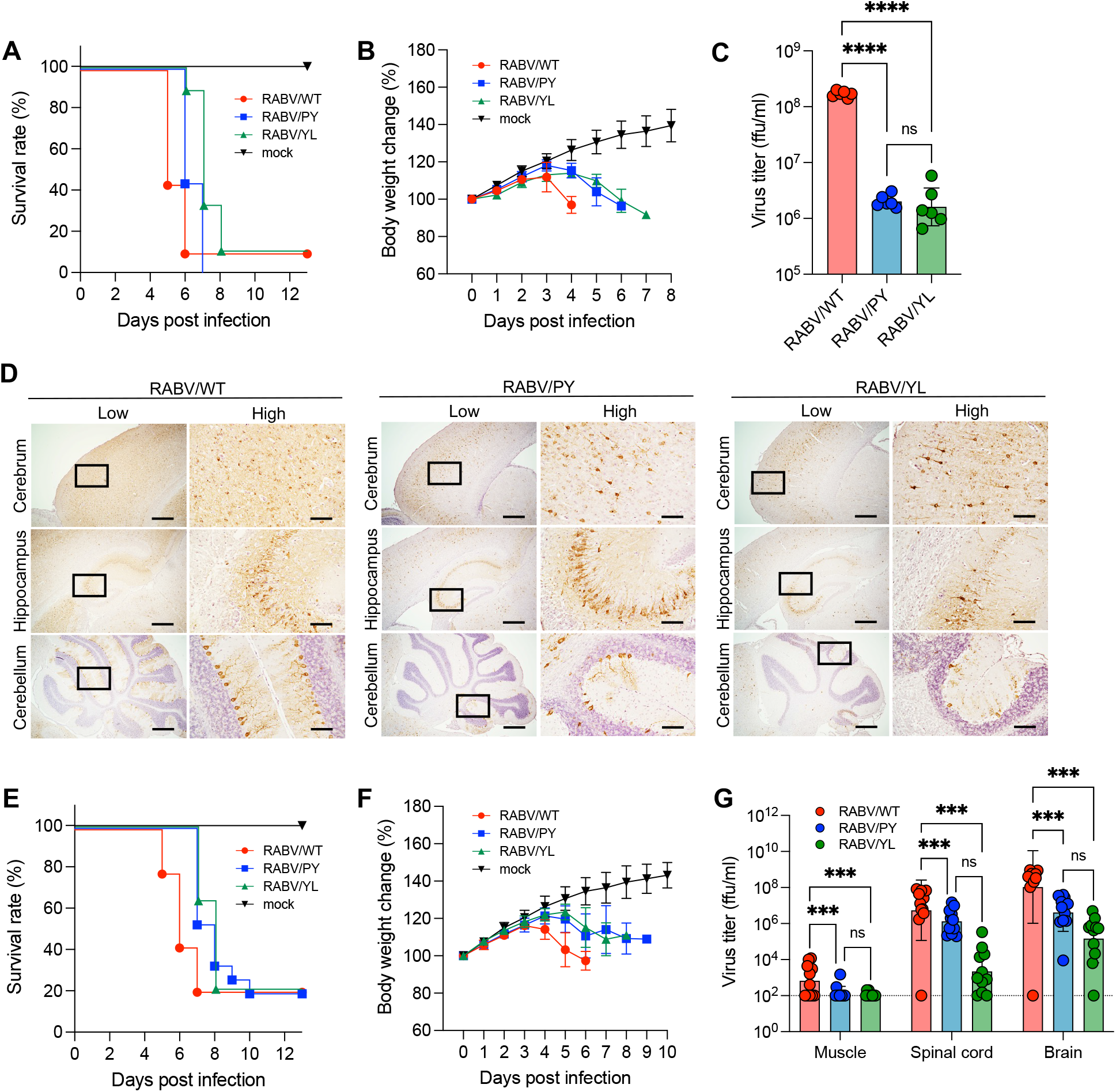
Attenuation of RABV L-domain mutants in vivo. Five-week-old ddY mice were inoculated with 10^2^ or 10^5^ ffu RABV intracranially (A–D) or intramuscularly (E–G). Virus-infected mice were monitored for (A, E) survival and (B, F) body weight changes. Data in the graphs are means ± standard deviations [mock: n = 6 (ic), n = 6 (im); RABV/WT: n = 8 (ic), n = 14 (im); RABV/PY: n = 9 (ic), n = 15 (im); RABV/YL: n = 9 (ic), n = 14 (im)]. (C, G) Virus titers in tissue homogenates at 4 dpi (ic) or 5 dpi (im) were determined using a focus forming assay. Bars show means ± standard deviations [n = 6 (ic) and n =11 (im) for each group]. (D) Immunohistochemistry of the mouse brain. Brain sections at 4 dpi (ic) were stained with anti-RABV N. Scale bars: 500 μm (low magnification) and 100 μm (high magnification). Statistical analyses: (C, G) one-way ANOVA and Tukey’s multiple *t*-tests: \*\*\**P* < 0.001, \*\*\*\**P* < 0.0001.

## Discussion

Virus budding is a sophisticated process required for infectious viral particle production and release. In rhabdovirus infection, the M protein is essential for budding and the formation of typical bullet-shaped particles, given that M-deficient viruses form long filamentous particles that are not infective^2^. RABV M also has multiple functions at the late stage of RABV infection, when it is involved in the condensation and assembly of RNPs as well as the pushing out and pinching off of viral particles^2,4–7,20^. However, little is known about the host factors involved in the budding and particle formation processes of rhabdoviruses. Our RNAi screen identified TSG101 as an ESCRT factor that supports RABV infection. Moreover, our experiments demonstrated that TSG101 facilitates RABV budding and virion formation.

The ESCRTs play roles in membrane scission, which is a crucial process in cytokinesis^9,11^ and the formation of endosomal luminal vesicles and exosomes^10^. ESCRT-mediated membrane invagination and vesicle formation is a process analogous to the budding step during the release of virions from cells. Some enveloped viruses, such as retroviruses, filoviruses, paramyxoviruses, and flaviviruses, are reported to use the ESCRT in viral particle budding^12,14,18,19^. In the present study, RABV growth was significantly impaired by TSG-KD (Fig. 1C); however, the knockdown of TSG101 had no effect on viral entry, genome replication, and protein synthesis (Fig. 2A–D), although it caused the intracellular accumulation of viral particles and the distortion of virion morphology (Fig. 2H–K). These results provide evidence that TSG101 contributes to the late stage of RABV infection, specifically virion assembly and budding.

L-domains are consensus amino acid sequences observed in the structural proteins of various viruses and are known to be involved in the recruitment of ESCRTs^21^. RABV possesses an L-domain composed of overlapping PY and YL motifs (PPEYVPL) at amino acid positions 38–41 in RABV M. Amino acid substitutions of the PY motif in RABV M are reported to decrease the growth and pathogenicity of RABV; however, the molecular mechanisms underlying this attenuation remain to be elucidated^16^. In the current study, we have demonstrated that TSG101 interacts with RABV M via the L-domain, which is required for the colocalization of these proteins (Figs. 3B and 4C). Recombinant RABVs lacking the L-domain in RABV M show low viral proliferation and pathogenicity both in vitro and in vivo (Fig. 5A; Fig. 6C, G). Notably, the RABV/YL mutant showed an impaired budding ability leading to an abnormal particle morphology (Fig. 5F). Meanwhile, the lack of PY motifs increased viral particle length, inhibited cell-to-cell infection, and reduced pathogenicity in mice (Fig. 5D-H, 6A-F), suggesting that this motif may contribute to RABV infection differently from the YL motif. Collectively, our results revealed an important function of the L-domain, especially the YL motif, in the TSG-dependent RABV life cycle. In contrast vesicular stomatitis virus, a prototype rhabdovirus, has been shown only to possess the PY motif in its M, and it is reportedly less dependent on TSG101 for viral growth^22^.

The binding mode of TSG101 to RABV M remains to be elucidated. In addition to constructing complexes with other ESCRTs, the key functions of TSG101 in endocytic trafficking are mediated by binding to one of the L-domains, i.e., to P(T/S)AP motif-containing proteins via methionine at position 95 (95M) or to ubiquitin via asparagine at position 45 (45N)^11,23–25^. None of the RABV proteins possess the P(T/S)AP motif, and our rescue experiments (Fig. 1F) and immunoprecipitation assays (Fig. 3C) suggest that 95M in TSG101 is not required for RABV replication. However, the introduction of an amino acid mutation on 45N (TSG/N45A) resulted in a poor RABV growth rescue rate (Fig. 1F) and a weakened RABV M binding capacity (Fig. 3C). Considering the role played by ubiquitin in triggering cell membrane invagination for endocytosis^26^, the involvement of ubiquitin in virus budding appears plausible^14,27–30^. During rhabdovirus infection, M is also ubiquitinated by the host E3 ubiquitin ligase^27,29^. Based on the present data and previous studies, we speculate that the ubiquitination of RABV M might facilitate the RABV M–TSG101 interaction.

In summary, this study has revealed that TSG101 supports the budding and formation of RABV viral particles through its interaction with RABV M via the L-domain. Considering the involvement of host factors in RABV infection, the discovery of a host factor that plays a critical role in the egress of RABV provides new insights into the mechanisms of RABV budding and virion formation, especially as it highlights the importance of the L-domain in RABV M. In future studies, focusing on the involvement of other ESCRT factors and ubiquitin will improve our understanding of the mechanisms underlying RABV budding and the production of bullet-shaped virion formation and may help integrate previously disconnected studies.

## Materials and Methods

### Ethics statement

Animal experiments were approved by the Institutional Animal Care and Use Committee of Hokkaido University (approval number 19-0014) and performed according to the committee’s guidelines.

### Cells

Human neuroblastoma (SK-N-SH) cells, mouse neuroblastoma (NA) cells, and baby hamster kidney cells stably expressing T7 RNA polymerase (BHK/T7-9) cells^31^ were maintained in Eagle’s Minimum Essential Medium (MEM) supplemented with 10% fetal bovine serum (FBS). SK-N-SH cells were cultured in type-I collagen–coated plates, whereas 293T cells were cultured in Dulbecco’s Modified Eagle’s Medium supplemented with 10% FBS. All cells were incubated at 37°C in the presence of 5% CO_2_.

SK-N-SH cells stably expressing TSG101–venus were established using lentivirus transduction. cDNA fragments encoding TSG101 fused at its N-terminus with venus following double GGGGS linker sequences were subcloned into the pLVSIN-CMV Pur vector. The resulting plasmid was transfected into 293T cells using Lentiviral High Titer Packaging Mix (Takara) to obtain the lentiviral vector for TSG101–venus transduction. The SK-N-SH cells were then infected with the lentivirus vector and selected using puromycin.

### Viruses

Recombinant RABV clones of the CVS strain, HEP strain, and a reporter CVS encoding NanoLuc [CVS-Nluc (N-P)] were prepared as described previously^32,33^. Recombinant RABV/PY and RABV/YL mutants were obtained from the cDNA clones of the CVS strain by substituting the PY and YL motif for alanine, respectively, which was achieved via PCR-based mutagenesis according to procedures described previously^31–33^. The street RABV Toyohashi strain was prepared as described previously^34^. All viruses were propagated in NA cells.

### Quantitative real-time RT-PCR

Viral RNA copy numbers were quantified using a Thunderbird Probe One-step qRT-PCR Kit (TOYOBO) and TaqMan probe/primer sets specifically targeting RABV CVS N (F: 5′-TCG AAT GCT GTC GGT CAT GT-3′; R: 5′-CCG AAG AAT TCC TCT CCC AAA TA-3′; probe: 5′-FAM-CAA TCT CAT TCA CTT TGT TG-MGB-3′).

### RNAi screen

A custom siRNA library targeting 25 ESCRT-related factors and consisting of three different siRNAs per target gene was used to rule out the off-target effect (Silencer Select predesigned siRNA, Ambion). SK-N-SH cells were reverse-transfected with 20 nM of siRNA using Lipofectamine RNAiMAX (Invitrogen) in collagen-coated 96-well plates and then cultured for 60 h. Subsequently, the cells were infected with CVS-Nluc (N-P) at a multiplicity of infection (MOI) of 10. At 18 hpi, the supernatants were transferred into NA cells seeded on 96-well clear-bottom black plates. After 8 h of incubation at 37L, luminescence signals were measured using a Nano-Glo Luciferase Assay (Promega) following the manufacturer’s protocol.

### Virus titration via a focus forming assay

NA cells were infected with serially diluted specimens in a 48-well plate for 1 h and then overlayed with MEM supplemented with 5% FBS, 0.5% methyl cellulose, and GlutaMAX (Gibco). After 3 days of incubation, the cells were fixed and stained with FITC-labelled anti-RABV N antibody (Fujirebio) as well as 10 μg/ml of Hoechst 33342. Foci were then counted to determine the virus titer in focus-forming units (ffu).

### Antiviral activity of ilaprazole, a TSG101 inhibitor

Cells were infected with viruses at a MOI of 1 in the presence of ilaprazole (SelleckChem) serially diluted in 5% FBS-supplemented MEM for 1 h. After the cells were washed with phosphate-buffered saline (PBS), fresh medium containing ilaprazole was added. Progeny viruses in the supernatants were titrated at 24 hpi.

### Rescue experiment

SK-N-SH cells were first transfected with 20 nM of siRNA against TSG101 using Lipofectamine RNAiMAX (Invitrogen). At 48 h post-transfection of siRNA, the cells were transfected with siRNA-resistant TSG101 constructs using Lipofectamine 2000 (Invitrogen). After 24 h of incubation, the cells were infected with CVS at a MOI of 1, and the supernatant was collected for virus titration at 24 hpi.

### Attachment and entry assay

The attachment and entry assays were conducted following procedures described previously^35^.

### Minigenome assay

A CVS strain-based minigenome assay was performed according to a previously reported method with some modification^36^ and with gene knockdown. Briefly, plasmids encoding CVS N, P, and L and the RABV minigenome were reverse-cotransfected into 293T cells in 96-well plates. After 8 h, the cells were transfected with 20 nM of siRNA using Lipofectamine RNAiMAX. Following 70 h of incubation, minigenome replication was evaluated by measuring the luminescence of NanoLuc from the minigenome using Nano-Glo (Promega).

### Immunofluorescence assays

RABV-infected cells were fixed with 10% phosphate-buffered formalin and washed with PBS. Blocking was conducted with 1% bovine serum albumin in PBS for 1 h at room temperature. The cells were then stained with anti-RABV M antibody (A54616; Epigentek; 1:150 dilution) in 0.3% BlockAce in 0.005% Tween-20 in PBS (PBST) overnight at 4°C. After two washes with PBST, Alexa Fluor 594–conjugated anti-rabbit IgG antibody (A32740; Invitrogen; 1:1000 dilution) was added to the cells as the secondary antibody. The stained cells were observed using a LSM780 confocal microscope and ZEN software (Zeiss).

### Electron microscopy analysis

For ultrathin-section electron microscopy, SK-N-SH cells in a collagen-coated dish (10 cm) were infected with rRABV at a MOI of 5–10 and cultured for 28 h. The cells were then washed with phosphate buffer (PB), scraped off the plate, and pelleted via centrifugation. The cell pellets were washed, fixed for 30 min with 2.5% glutaraldehyde in PB on ice, and fixed again with freshly prepared fixative at 4°C overnight. The fixed cell pellets were washed with PB and postfixed with 1% osmium tetroxide in PB for 1 h on ice. Subsequent procedures were conducted as described previously^37^.

For negative-stain electron microscopy, rRABV purified via ultracentrifugation with a 20% sucrose cushion was fixed with 2.5% glutaraldehyde. The fixed virus was placed on a elastic carbon film ELS-C10 (10-1012, Oken) and stained with 2% phosphotungstic acid solution (pH 5.8). The samples were then examined using a H-7650 electron microscope (Hitachi).

### Immunoprecipitation assay

First, 293T cells were cotransfected with plasmids encoding FLAG- or HA-tagged TSG101 and RABV M using polyethylenimine. Cell lysates were harvested at 72 h post-transfection, clarified via centrifugation. Immunoprecipitation was then conducted using anti-HA magnetic beads (Pierce) according to manufacturer’s protocol. The precipitates were extracted into 1x SDS sample buffer containing 2-mercaptethanol and subjected to a standard immunoblotting assay using HRP-conjugated anti-HA antibody (H6533; Sigma; 1:3000 dilution) and anti-FLAG antibody (A8592; Sigma; 1:3000 dilution).

### Animal experiments

Five-week-old male ddY mice were used in the animal experiments. Mice were inoculated intracranially or intramuscularly with 10^2^ or 10^5^ ffu of rRABV under anesthesia using isoflurane. All mice in the survival groups were observed daily for disease symptoms and bodyweight changes. The humane endpoint was defined as a 20% decrease in body weight or an inability to reach food or water due to the onset of disease. Mice were euthanized by decapitation under anesthesia, and their tissues were harvested at specific time points for further analysis.

### Immunohistochemistry

Brain sections were prepared as described previously^32^. After antigen retrieval in citrate buffer for 5 min using a pressure cooker, the sections were treated with 0.3% H_2_O_2_ in methanol for 15 min to inactivate endogenous peroxidase. The sections were then treated with 10% goat serum (Nichirei Biosciences) for 1 h at room temperature and incubated at 4L overnight with the primary antibody against RABV N (PA321352LA01RAI; Cusabio; 1:2500 dilution). After three washes with 0.01% PBST, secondary staining was performed with EnVision system-HRP Labelled Polymer Anti-Rabbit (K4220, Dako) for 1 h at room temperature. The slides were washed three times with PBST, and a DAB Substrate Kit (425011, Nichirei) was used to visualize the immunostaining.

### Statistical analysis

All statistical analyses were performed using GraphPad Prism 9.2.0. For comparisons of two groups, Welch’s *t*-test was used. For comparisons of two groups at multiple time points, a multiple *t*-test was performed using the Benjamini, Krieger, and Yekutieli method. For comparisons among more than two groups, one-way/two-way ANOVA with Tukey’s/Dunnett’s multiple comparisons test was used. Data are presented as means ± standard deviations in graphs. \**P* < 0.05, \*\**P* < 0.01, \*\*\**P* < 0.001, \*\*\*\**P* < 0.0001.

## Data availability

This study includes no data deposited in external repositories.

## Acknowledgments

Our great appreciation is extended to Mr. T. Namba, Dr. O. Ichii, and Prof. Y. Kon for supporting for uranyl acetate staining in the ultrathin-section electron microscopic analyses.

This study was supported in part by the Japan Society for the Promotion of Science (JSPS) KAKENHI under grant numbers 18H02333, 18KK0192, 21J13419; the Japan Agency for Medical Research and Development (AMED) under grant number JP22fk0108141; Health Labour Sciences Research under grant number 22HA0601; and the World-leading Innovative and Smart Education (WISE) Program from the Ministry of Education, Culture, Sports, Science, and Technology (MEXT) (1801).

## Author Contributions

Y.I. and M.S. designed the research; Y.I., K.T., T.S., K.I., N.K., M.K., S.T., S.K., M.H., M.S. and Y.O. performed experiments and analyzed data; S.I., K.M., N.I., A.T., W.H., Y.O. and H.S. provided critical reagents and advice; Y.I. and M.S. wrote the draft manuscript; M.S., N.I., A.T., W.H., Y.O., and H.S. provided critical reviews of certain experiments and of the draft manuscript; M.S., N.I., K.M., H.S. and Y.I. acquired funding support.

## References

1. Finke S. & Conzelmann KK. Replication strategies of rabies virus. Virus Res. 111, 120–131 (2005).

2. Mebatsion T., Weiland F. & Conzelmann. KK. Matrix protein of rabies virus is responsible for the assembly and budding of bullet-shaped particles and interacts with the transmembrane spike glycoprotein G. J. Virol. 73, 242–250 (1999).

3. Cadd T. L., Skoging U. & Liljeström P. Budding of enveloped viruses from the plasma membrane. Bioessays. 19, 993–1000 (1997).

4. Kaptur P. E., Rhodes R. B. & Lyles D. S. Sequences of the vesicular stomatitis virus matrix protein involved in binding to nucleocapsids. J. Virol. 65, 1057–1065 (1991).

5. Lyles D. S. & McKenzie M. O. Reversible and irreversible steps in assembly and disassembly of vesicular stomatitis virus: equilibria and kinetics of dissociation of nucleocapsid-M protein complexes assembled in vivo. Biochemistry 37, 439–450 (1997).

6. Newcomb W. W. & Brown J. C. Role of the vesicular stomatitis virus matrix protein in maintaining the viral nucleocapsid in the condensed form found in native virions. J. Virol. 39, 295–299 (1981).

7. Lyles D. S., McKenzie M. & Parce J. W. Subunit interactions of vesicular stomatitis virus envelope glycoprotein stabilized by binding to viral matrix protein. J. Virol. 66, 349–358 (1992).

8. Mebatsion T., König M. & Conzelmann KK. Budding of rabies virus particles in the absence of the spike glycoprotein. Cell. 84, 941–951 (1996).

9. Hurley J. H. & Hanson P. I. Membrane budding and scission by the ESCRT machinery: It’s all in the neck. Nat Rev Mol Cell Biol. 11, 556–566 (2010).

10. Colombo M. et al. Analysis of ESCRT functions in exosome biogenesis, composition and secretion highlights the heterogeneity of extracellular vesicles. J Cell Sci. 126, 5553–5565 (2013).

11. Hurley J. H. & Emr S. D. The ESCRT complexes: Structure and mechanism of a membrane-trafficking network. Annu Rev Biophys Biomol Struct. 35, 277–298 (2006).

12. Garrus J. E. et al. Tsg101 and the vacuolar protein sorting pathway are essential for HIV-1 budding. Cell 107, 55–65 (2001).

13. Urata S. et al. Interaction of TSG101 with Marburg virus VP40 depends on the PPPY motif, but not the PT/SAP motif as in the cse of Ebola virus, and TSG101 plays a critical role in the budding of Marburg virus-like particles induced by VP40, NP, and GP. J Virol. 81, 4895–4899 (2007).

14. Timmins J. et al. Ebola virus matrix protein VP40 interaction with human cellular factors TSG101 and Nedd4. J Mol Biol. 326, 493–502 (2003).

15. Zhang Y. et al. Proteomic Profiling of Purified Rabies Virus Particles. Virol Sin. 35, 143–155 (2020).

16. Wirblich C. et al. PPEY motif within the rabies virus (RV) matrix protein. is essential for efficient virion release and RV pathogenicity. J Virol. 82, 9730–9738 (2008).

17. Leis J., Luan C. H., Audia J. E., Dunne S. F., & Heath C. M. Ilaprazole and other novel prazole-based compounds that bind TSG101 inhibit viral budding of Herpes Simplex Virus 1 and 2 and Human Immunodeficiency Virus from Cells. J Virol. 95, e00190–21 (2021).

18. Park A. et al. Nipah Virus C Protein recruits TSG101 to promote the efficient release of virus in an ESCRT-dependent pathway. PLoS Pathog. 12, e1005659 (2016).

19. Tabata K. et al. Unique requirement for ESCRT factors in flavivirus particle formation on the endoplasmic reticulum. Cell Rep. 16, 2339–2347 (2016).

20. Justice P. A. et al. Membrane vesiculation function and exocytosis of wild-type and mutant matrix proteins of vesicular stomatitis virus. J. Virol. 69, 3156–3160 (1995).

21. Chen B. J. & Lamb R. A. Mechanisms for enveloped virus budding: Can some viruses do without an ESCRT? Virology. 372, 221–232 (2008).

22. Irie T., Licata J. M., McGettigan J. P., Schnell M. J. & Harty R. N. Budding of PPxY-containing rhabdoviruses is not dependent on host proteins TGS101 and VPS4A. J Virol. 78, 2657–2665 (2004).

23. Pornillos O. et al. Structure and functional interactions of the TSG101 UEV domain. EMBO J. 21, 2397–2406 (2002).

24. Nickerson D. P., Russell M. R. G. & Odorizzi G. A concentric circle model of multivesicular body cargo sorting. EMBO Rep. 8, 644–650 (2007).

25. Ferraiuolo R. M., Manthey K. C., Stanton M. J., Triplett A. A. & Wagner K. U. The multifaceted roles of the tumor susceptibility gene 101 (TSG101) in normal development and disease. Cancers 12, (2020).

26. Strous G. J. & Govers R. The ubiquitin-proteasome system and endocytosis. J Cell Sci. 112, 1417–1423 (1999).

27. Harty R. N., Paragas J., Sudol M. & Palese P. A proline-rich motif within the matrix protein of vesicular stomatitis virus and rabies virus interacts with WW domains of cellular proteins: implications for viral budding. J. Virol. 73, 2921–2929 (1999).

28. Sette P., Jadwin J. A., Dussupt V., Bello N. F. & Bouamr F. The ESCRT-associated protein Alix recruits the ubiquitin ligase Nedd4-1 to facilitate HIV-1 release through the LYPX_n_L L domain motif. J Virol. 84, 8181–8192 (2010).

29. Harty R. N. et al. Rhabdoviruses and the cellular ubiquitin-proteasome system: a budding interaction. J Virol. 75, 10623–10629 (2001).

30. Yasuda J., Nakao M., Kawaoka Y. & Shida H. Nedd4 regulates egress of Ebola virus-like particles from host cells. J Virol. 77, 9987–9992 (2003).

31. Ito N. et al. Improved recovery of rabies virus from cloned cDNA using a vaccinia virus-free reverse genetics system. Microbiol. Immunol. 47, 613–617 (2003).

32. Itakura Y. et al. Glu_333_ in rabies virus glycoprotein is involved in virus attenuation through astrocyte infection and interferon responses. iScience. 25, (2022).

33. Anindita P. D. et al. Generation of recombinant rabies viruses encoding NanoLuc luciferase for antiviral activity assays. Virus Res. 215, 121–128 (2016).

34. Nosaki Y. et al. Fourth imported rabies case since the eradication of rabies in Japan in 1957. J Travel Med. 28, 1–4 (2021).

35. Sabino C., Bender D., Herrlein M. L. & Hildt E. The epidermal growth factor receptor is a relevant host factor in the early stages of the Zika virus life cycle in vitro. J. Virol. 95, e01195–21 (2021).

36. Anindita P. D. et al. Ribavirin-related compounds exert in vitro inhibitory effects toward rabies virus. Antiviral Res. 154, 1–9 (2018).

37. Noda T. et al. Ebola virus VP40 drives the formation of virus-like filamentous particles along with GP. J Virol. 76, 4855–4865 (2002).

