## Supplemental Figures for "Morphogenesis of bullet-shaped rabies virus particles requires a functional interplay between the viral matrix protein and ESCRT-I component TSG101"

**Supplemental Information**

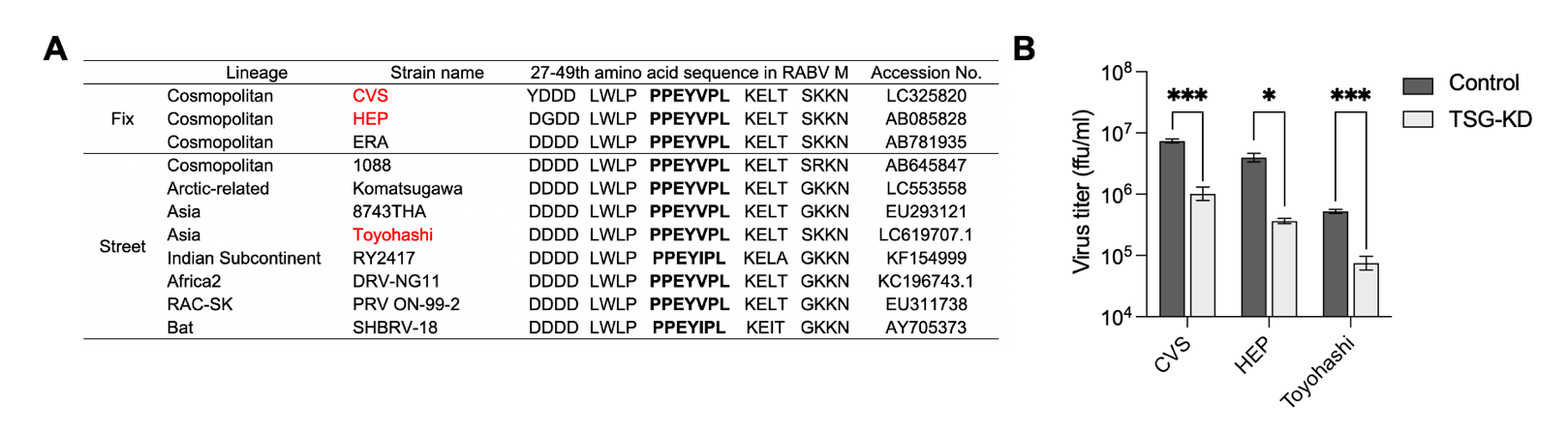

**Fig. S1. Conservation and functional importance of the L-domain in various fixed and street strains of RABV (related to Fig. 3).**

(A) Table listing the 27^th^ to 49^th^ amino acid sequences in the RABV M. Bold: L-domain; red: strains analyzed in (B). (B) Virus titers of the CVS strain, HEP fixed strain, and Toyohashi street strain in TSG-KD cells. SK-N-SH cells were infected with RABV at a MOI of 1. Virus titers in the culture supernatants at 48 hpi were determined using a focus forming assay. Bars indicate the means ± standard deviations of three replicates from a representative experiment. Statistical analyses: multiple *t*-tests with Welch’s correction: **P* < 0.05, ****P* < 0.001.

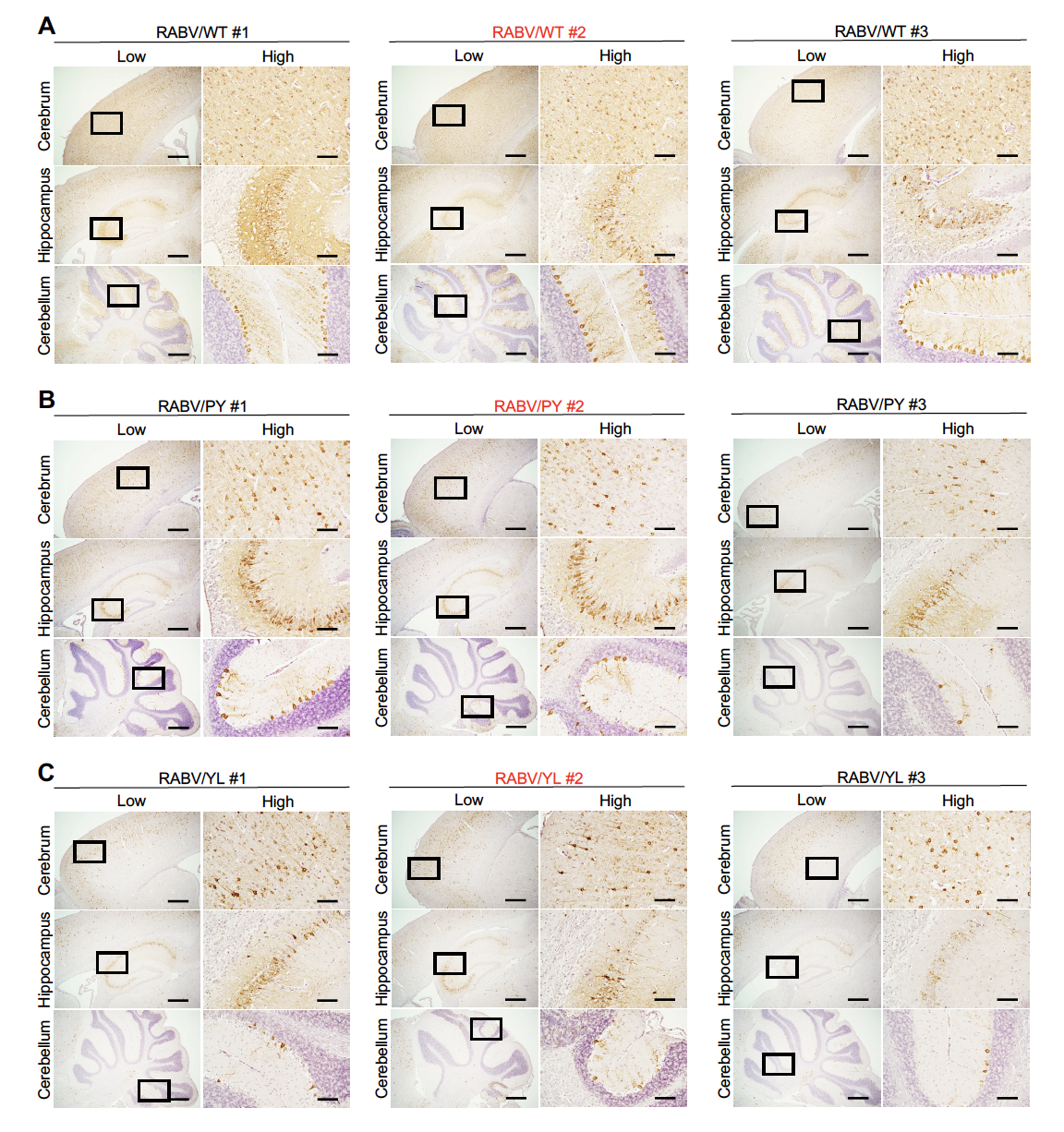

**Fig. S2. Distribution of RABV in mouse brains on 4 dpi (related to Fig.6).**

Immunohistochemistry of the mouse brain. Five-week-old ddY mice were inoculated with 10^2^ ffu RABV intracranially. Brain sections at 6 dpi of (A) RABV/WT, (C) RABV/PY, or (D) RABV/YL were stained with anti-RABV N. Scale bars: 500 μm (low magnification) and 100 μm (high magnification). Red-highlighted data are shown in the main figure as representative images.

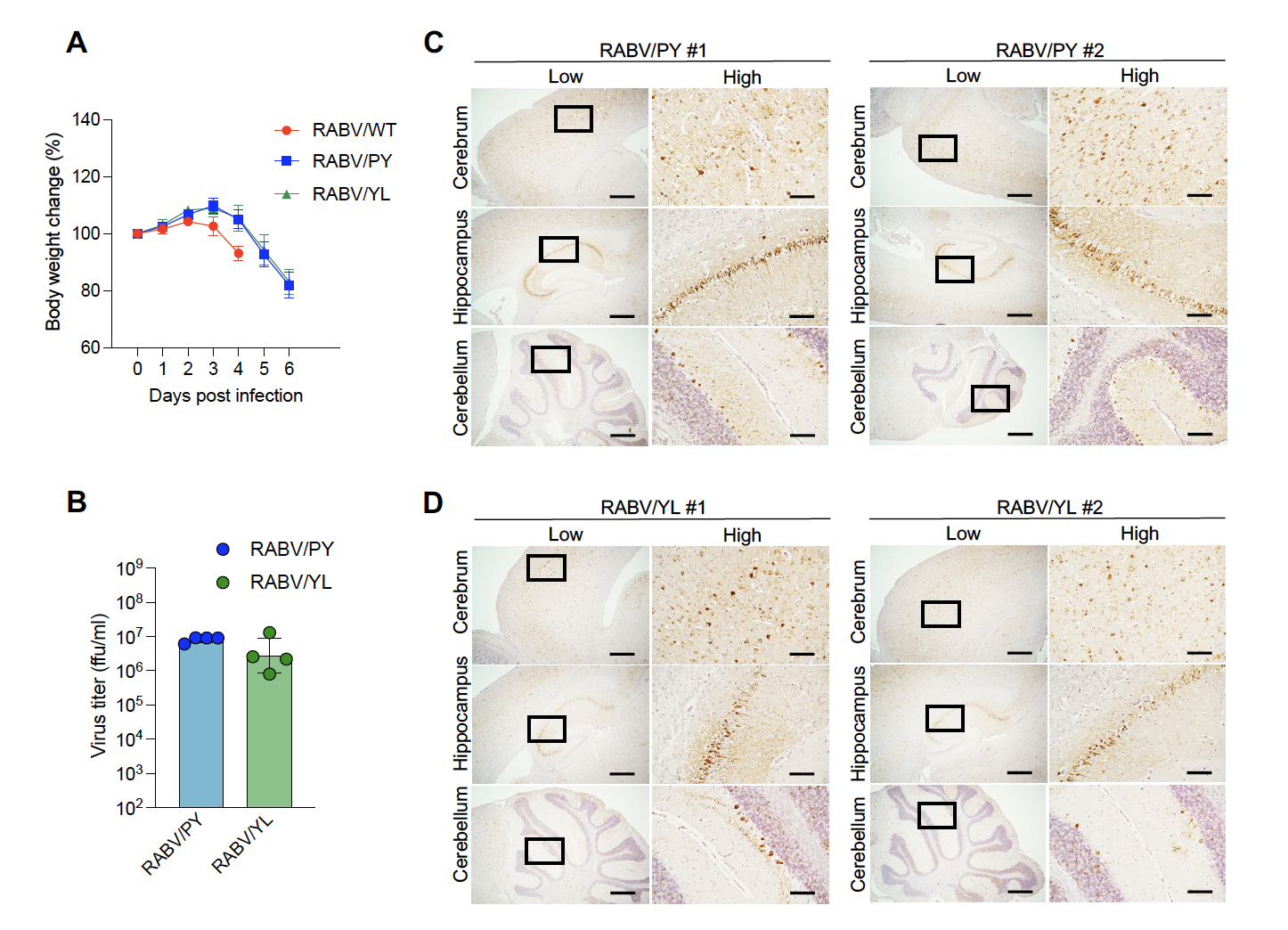

**Fig. S3. Virus growth and distributions of RABV mutants in mouse brains on 6 dpi (related to Fig.6).**

Five-week-old ddY mice were inoculated with 10^2^ ffu RABV intracranially. Virus-infected mice were monitored for (A) body weight changes. Data in the graphs are means ± standard deviations (RABV/WT: n = 3; RABV/PY: n = 6; RABV/YL: n = 6). (B) Virus titers in brain homogenate at 6 dpi were determined using a focus forming assay. Bars show means ± standard deviations (n = 6). (C, D) Immunohistochemistry of the mouse brain. Brain sections at 6 dpi by (C) RABV/PY or (D) RABV/YL were stained with anti-RABV N. Scale bars: 500 μm (low magnification) and 100 μm (high magnification).
